# Intraline genomic heterogeneity of the triple-negative breast cancer MDA-MB-231 cell line

**DOI:** 10.1101/2025.02.13.638020

**Authors:** Nair Varela-Rouco, Nuria Estévez-Gómez, Cristóbal Fernández-Santiago, Laura Tomás, Miriam Pérez, Daniel García-Souto, Juan Jose Pasantes, Roberto Piñeiro, João Miguel Alves, David Posada

## Abstract

Cancer cell lines are valuable models for studying tumor biology, yet their genomic evolution during culture can compromise experimental reproducibility. We conducted a detailed genomic analysis of the triple-negative breast cancer cell line MDA-MB-231, examining sublines obtained from different sources, at various time points, and across distinct passages. We introduce the concept of intraline heterogeneity (ILH) to highlight the genomic variability observed among these sublines. Our analyses revealed extensive genomic diversity, including differences in single nucleotide variants (SNVs) and copy number alterations (CNAs). In particular, CNAs exhibited remarkable heterogeneity, with pronounced chromosomal gains and losses between sublines, underscoring the impact of genomic instability on ILH. These findings suggest that ILH may influence experimental outcomes, emphasizing the importance of considering passage-specific genomic characterization to ensure consistency and reliability in cancer research.

## INTRODUCTION

Cancer cell lines derived from human tumors are widely utilized in cancer research due to their ease of use and ability to provide an unlimited source of biological material while retaining key genetic features of the original tumor ^1–5^. However, cell lines are not genetically stable; they accumulate mutations over generations and through culture passages, leading to significant genomic evolution ^6–8^. For example, a comprehensive genomic analysis of 27 passages of the breast cancer cell line MCF7 revealed rapid genetic diversification among passages which led to significant differences in drug response ^9^. Similar studies have additionally shown that chromosomal stability can be compromised due to sequential selection during long-term culture, leading to substantial karyotypic differences across passages ^10^. Although initiatives such as the Cancer Cell Line Encyclopedia (CCLE) ^11^ and the Catalogue of Somatic Mutations in Cancer (COSMIC) cell lines project ^12,13^ provide a genomic description of widely used cancer cell lines, these catalogs do not typically account for genomic changes acquired during culture passages. This unrecognized variability may lead to misinterpretation of experimental results, such as misidentifying passage-specific mutations as *de novo* mutations within xenograft experiments. Therefore, understanding cell line genomic heterogeneity is critical, as it can influence experimental reproducibility and the biological behavior of cell lines, potentially affecting the validity of preclinical findings and derived therapeutic strategies.

Here we present a comprehensive genomic characterization of the MDA-MB-231 cell line, a highly aggressive, invasive, and poorly differentiated triple-negative breast cancer (TNBC) cell line, established in 1973 from a pleural effusion of a 51-year-old Caucasian woman with metastatic breast cancer ^14^. This cell line is widely used as a model of advanced and invasive breast cancer due to its inherent ability to metastasize spontaneously to lymph nodes in xenograft experiments ^15^. Its intrinsic molecular subtype classification is under debate, shifting from basal-like ^16,17^ to claudin-low ^18,19^. Cytogenetic studies have revealed significant karyotypic heterogeneity, including the absence of chromosome 8 in most metaphases, chromosome counts ranging from 57 to 68 with a near-triploid copy number, and some unassignable chromosomes or chromosome parts ^20,21^.

To better understand the levels of genomic heterogeneity in MDA-MB-231, we sequenced multiple acquisitions and passages of this cell line. We introduce the term *intraline heterogeneity* (ILH) to describe the genomic variation observed among distinct passages of this cell line. This variation can arise regardless of whether they refer to passages produced within the same laboratory or purchases made at different time points or from distinct suppliers. Our analysis demonstrates extensive ILH in MDA-MB-231, highlighting its potential to influence experimental outcomes and emphasizing the importance of accounting for passage-specific genomic variability in cancer research.

## RESULTS

We characterized three commercial batches (CL0, CL1, and CL2) of the MDA-MB-231 cell line. CL0 has been maintained in the laboratory for over seven years, while CL1 and CL2 were acquired four and two years ago, respectively. Over the past four years, CL1 underwent five culture passages (CL1.E01, CL1.E02, CL1.E03, CL1.E04, and CL1.E06), with passage numbers not necessarily reflecting chronological order.

### The karyotype analysis of MDA-MB-231 confirmed near-triploidy

Our cytogenetic survey of 42 MDA-MB-231 (CL1) metaphases revealed an average chromosome count of 62.12, with a median of 60.5 and a mode of 63 chromosomes, within a range of 57 to 87 (Figure 1). The first and third quartiles were 59 and 63 chromosomes, respectively. We also identified sporadic double-minute chromosomes in several metaphases. These observations suggest a chromosomal count consistent with near-triploidy in agreement with that reported by Satya-Prakash *et al*. ^20^, who found a modal chromosome number of 57 within a range of 52 to 62, as well as data from American Type Culture Collection (ATCC), which reported a modal number of 64 and a range of 52 to 68 for this cell line. The karyotype reconstruction based on DAPI banding patterns, equivalent to G-banding but accentuating heterochromatic regions of specific chromosomes ^22^, shares many of the characteristics previously described for this cell line ^20,21^.

**Figure 1.**
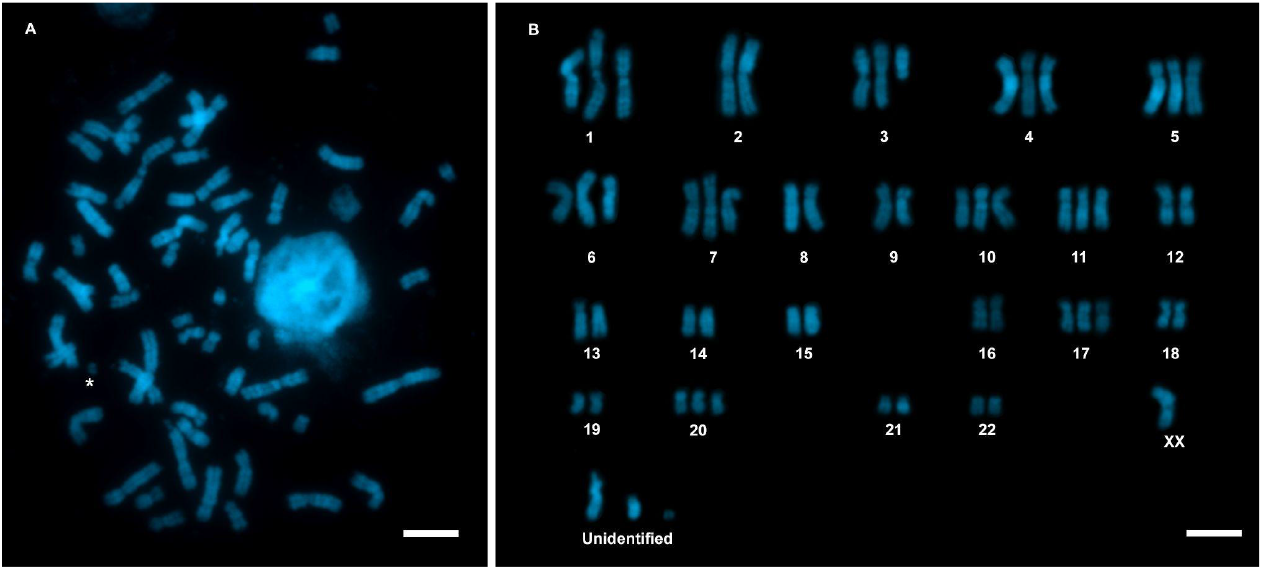
Metaphase spread and karyotype of the MDA-MB-231 (CL1) cell line. (A) Representative metaphase spread. The asterisk (*) marks the presence of a double minute chromosome. (B) Karyotype representation. Chromosomes are labeled according to their corresponding pairs. (A-B) The scale bar represents 10 µm.

### High levels of copy number heterogeneity among sublines

The copy number (CN) analysis revealed widespread chromosomal aneuploidies across the genome (Figure 2A), with significant heterogeneity in gains and losses among sublines (CL0, CL1, and CL2) and, to a lesser extent, within CL1 passages (E01–E06). We detected conserved diploid regions in whole chromosomes (13, 15, 16, 21, and 22) and parts of chromosomes (1, 2, 6, and 8). Gains were consistently observed in regions of chromosomes 1, 4, 5, 9, 10, 11, 14, 17, and 19. Deletions were identified in specific regions of chromosomes 9 (all sublines), 18 (CL1 and CL2), and 1 (CL1 passages E01, E02, and E06). The analysis at the level of breast cancer (BRCA) driver genes further confirmed the CN heterogeneity within sublines and passages (Figure 2B). All sublines exhibited copy number gains (CN=3) in *NOTCH1, NOTCH2, PTEN, FOXA1, TP53, ERBB2, BRCA1, USP6*, and *BRIP1*; Additional gains were observed in *PIK3CA, NF1*, and *MAP3K13* (CN=3 for CL1 and CL2; CN=4 for CL0) as well as *FAT3* and *ASXL1* (CN=4 across all sublines). Conversely, copy number losses (CN=1) were detected in *JAK2* and *PTPRD* in all sublines. Subline CL0 displayed unique differences compared to CL1 and CL2: gains of CN=3 in *DNMT3A, ALK, BIRC6*, and *POLQ* (CN=2 in CL1 and CL2), CN=4 in *PIK3CA, MAP3K13*, and *NF1* (CN=3 in CL1 and CL2); and CN ≥ 5 in *TBX3*, and *CSMD3* (CN=3–4 in CL1 and CL2). Furthermore, CL0 retained diploid copy numbers (CN=2) in genes such as *PBRM1, FAT4, FAT1, GATA3, IRS4, ERBB3*, and *KMT2D*, (CN=3–4 for CL1 and CL2), as well as CN=2 for *SMAD4* (CN=1 in CL1 and CL2).

**Figure 2.**
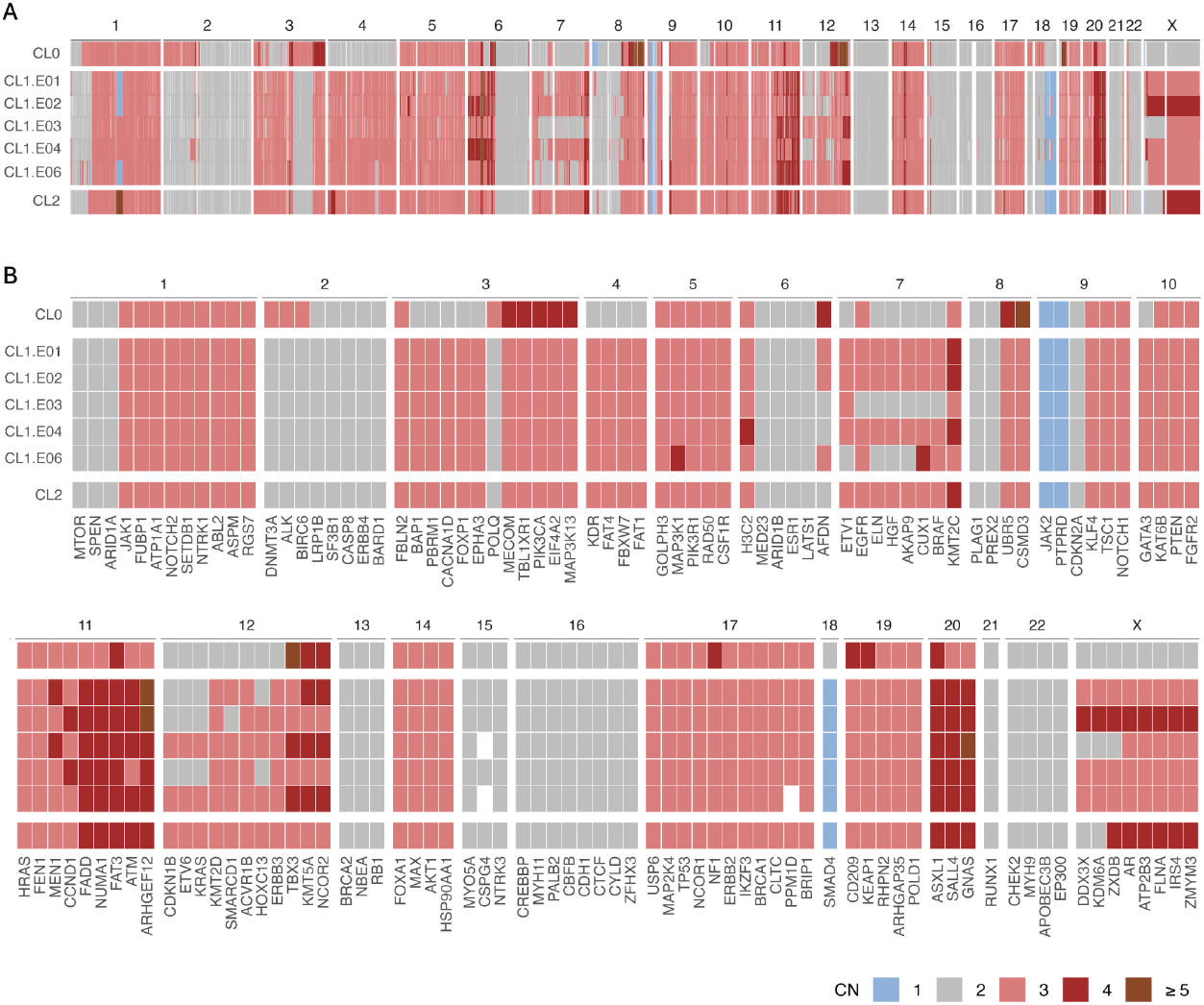
Copy number profiles of all MDA-MB-231 sublines. (A) Genome-wide copy number (CN). Different facets reflect chromosomes (columns sorted by genomic position) and samples (rows). Colors indicate CN status. CN ≥ 5 was collapsed into a single category. (B) CN per BRCA driver gene is ordered according to chromosome location (columns, genes are sorted by genomic position within each chromosome) and samples (rows).

### Genomic differences among MDA-MB-231 sublines based on SNVs

The observed sequencing depth (293–1,096 X) and breadth (>99.6%) were sufficient for a comprehensive genomic characterization of all sublines (Table S1). Across all sublines, we identified 28,073 mutations —26,173 single nucleotide variants (SNVs) and 1,900 small insertions and deletions (indels)— (Figure 3). Notably, 26,193 mutations (24,721 SNVs and 1,472 indels) were shared among all sublines. CL1 and CL2 shared 1,012 mutations (961 SNVs and 51 indels) with a mean variant allele frequency (VAF) < 0.5. Subline CL0 harbored 192 private mutations at variable frequencies, with some exceeding 0.5. CL2 had 26 private mutations with a mean VAF < 0.5. Within CL1, CL1.E03 shared the fewest mutations with the remaining passages despite having the highest sequencing depth (Table S1). The VAF distribution revealed a complex ploidy pattern across chromosomes (Figure 4). For example, chromosomes 2 and 21 displayed clear VAF peaks at 0.5, indicating diploidy. In contrast, chromosomes 5, 9, 10, and 20 exhibited peaks at 0.33 and 0.66, consistent with triploidy, whereas chromosome 18 in CL1 and CL2 or chromosome 3 in CL0 showed peaks around 0.25 and 0.75, suggesting tetraploidy. Overall, the VAF distributions per chromosome significantly differed among sublines, with the largest difference observed between CL0 and the other two sublines.

**Figure 3.**
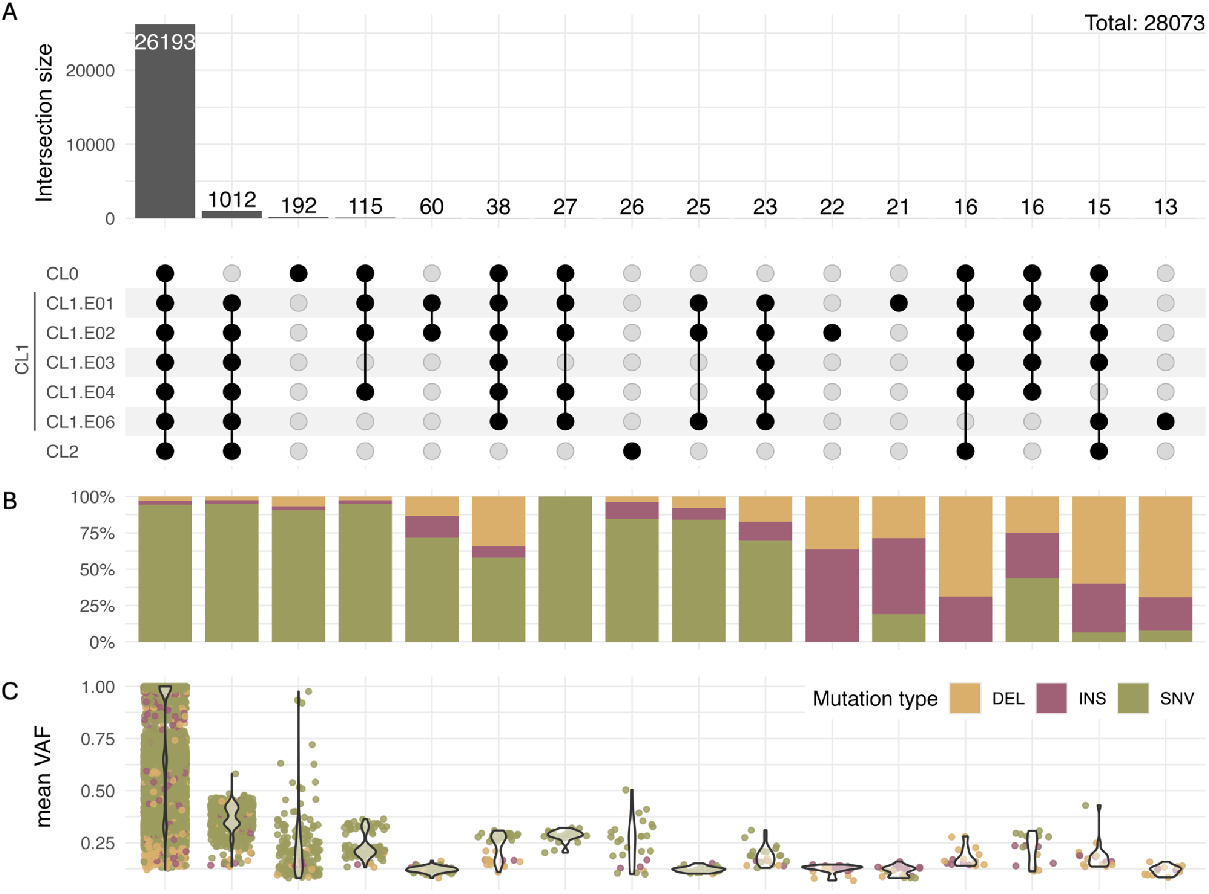
Mutational landscape and genomic heterogeneity of MDA-MB-231. (A) Upset plot depicting the number of mutations (SNVs + indels) shared across samples (intersections). Bars represent the number of mutations at each intersection, while black dots below indicate which samples are involved in each intersection. Only intersections with more than 12 mutations are shown. (B) Mutation type distribution, showing the percentage of deletions (DEL), insertions (INS), and single nucleotide variants (SNVs) for each intersection. (C) Mean variant allele frequency (VAF) distribution, colored by mutation type.

**Figure 4.**
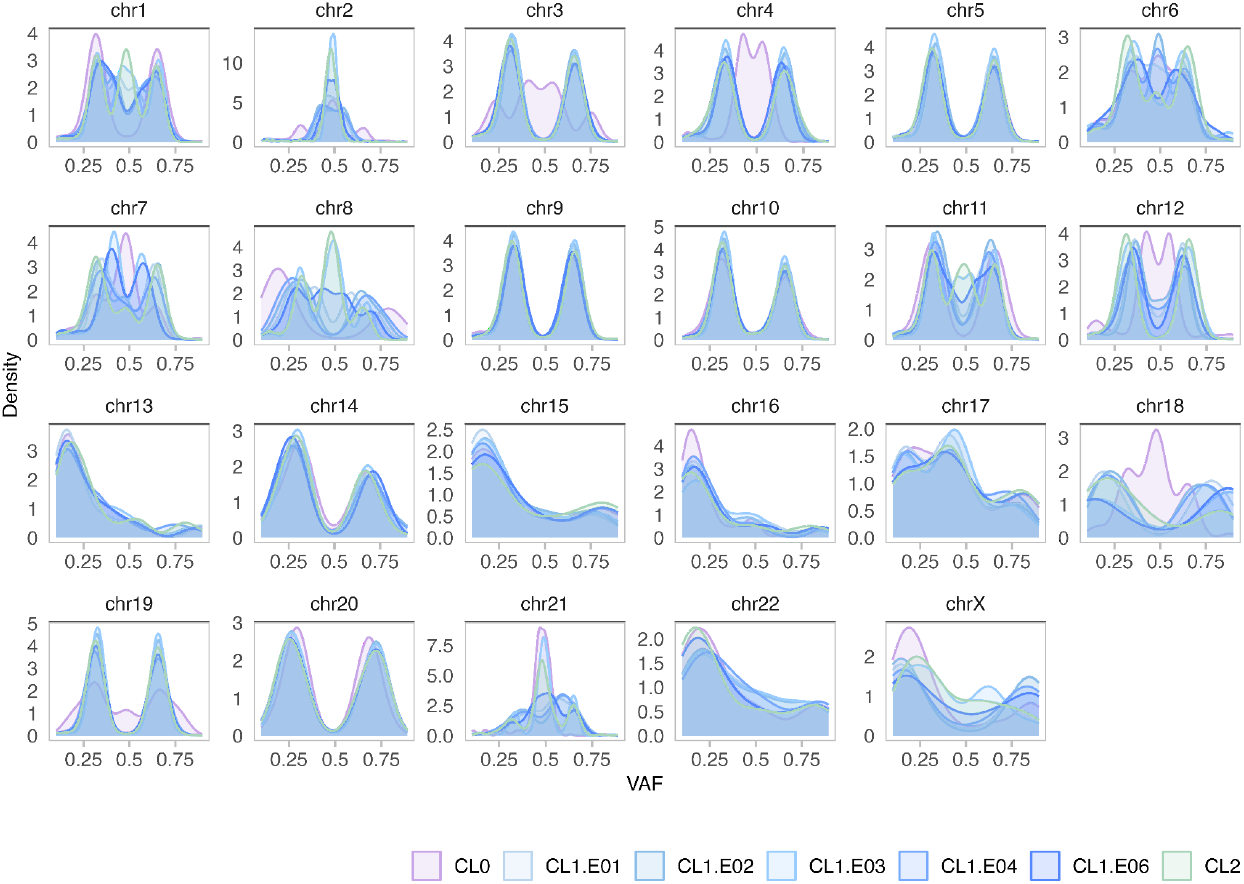
Chromosome-wide variant allele frequency distribution across sublines. VAF density distribution of SNVs for each chromosome and each sample. For clarity, only VAF values ranging from 0.1 to 0.9 are shown. Different colors represent distinct sublines: CL0, CL1 (E01, E02, E03, E04, and E06), and CL2. Within CL1, distinct shades of blue indicate different passages.

### Non-synonymous SNVs VAF variations across sublines in BRCA driver genes

We next explored mutations affecting BRCA driver genes reported in the COSMIC ^23^ and IntoGen ^24^ databases. We identified 38 BRCA drivers harboring 78 non-synonymous SNVs, all of which were present in all sublines (Figure 5, Table S2). Of these, eight —*BRAF*-G464V, *FAT3*-G1873V, *FAT3*-S4319F, *KRAS*-G13D, *ASPM*-S1422P, *LRP1B*-R191I, *PBRM1*-I228V, and *TP53*-R280K— have already been described for the MDA-MD-231 cell line in the COSMIC Cell Lines Project v.101 ^13^. We further investigated ten SNVs in six driver genes (*AKAP9, BRAF, CDKN1B, ELN, KRAS*, and *SPEN*) that exhibited VAF differences among sublines (standard deviation > 0.1) (Table 1). Subline CL0 displayed the most pronounced VAF variation compared to CL1 (all passages) and CL2. Specifically, CL0 showed increased VAFs for *AKAP9*-N2792S, *AKAP9*-M463I, *AKAP9*-M3614V, *BRAF*-G464V, *CDKN1B*-V109G, *KRAS*-G13D, and *SPEN*-R2283K, and decreased VAFs for *ELN*-G422S, *SPEN*-L1091P, and *SPEN*-N2360D in contrast with CL1 and CL2. Among these, two SNVs (*BRAF*-G464V and *KRAS*-G13D) were previously reported for the MDA-MB-231 cell line ^13,25,26^. Seven (*AKAP9*-N2792S, *AKAP9*-M463I, *AKAP9*-M3614V, *CDKN1B*-V109G, *ELN*-G422S, *SPEN*-L1091P, and *SPEN*-N2360D) are described in ClinVar ^27^ and one (*SPEN*-R2283K) has not been previously reported. From the seven SNVs reported in ClinVar, *AKAP9*-N2792S, and *AKAP9*-M463I have been associated with breast cancer risk ^28,29^, while *CDKN1B*-V109G was found in non-TNBC-samples ^30–33^. Interestingly, some of the SNVs we detected have been associated with other diseases: *AKAP9*-M3614V has been reported in cardiac disease ^34,35^; *ELN*-G422S has been identified in gastrointestinal cancers ^13^; and both *SPEN*-L1091P and *SPEN*-N2360D are described in ClinVar as associated with Radio-Tartaglia syndrome and *SPEN*-related disorders ^27^.

**Table 1.** Non-synonymous SNVs with VAF differences among sublines. Gene symbol, genomic position for the hg38 reference, amino acid change code, germline status in ClinVar ^27^, and whether it was reported for MDA-MB-231 in COSMIC with the verified status in parenthesis ^13^.

| Gene symbol | Genomic position for hg38 | Amino acid change | Germline status in ClinVar | Reported for MDA-MB-231 |
| --- | --- | --- | --- | --- |
| **AKAP9** | chr7:92083384.A>G | N2792S | Benign | - |
|  | chr7:92001306.G>T | M463I | Benign | - |
|  | chr7:92099813.A>G | M3614V | Benign | - |
| **BRAF** | chr7:140781617.C>A | G464V | Pathogenic / likely pathogenic | Yes (verified) |
| **CDKN1B** | chr12:12718165.T>G | V109G | Benign / uncertain | - |
| **ELN** | chr7:74056384.G>A | G422S | Benign / likely benign | - |
| **KRAS** | chr12:25245347.C>T | G13D | Pathogenic | Yes (verified) |
| **SPEN** | chr1:15929512.T>C | L1091P | Benign | - |
|  | chr1:15933318.A>G | N2360D | Benign | - |
|  | chr1:15933088.G>A | R2283K<sup>a</sup> | - | - |

<sup>a</sup> Not previously reported.

**Figure 5.**
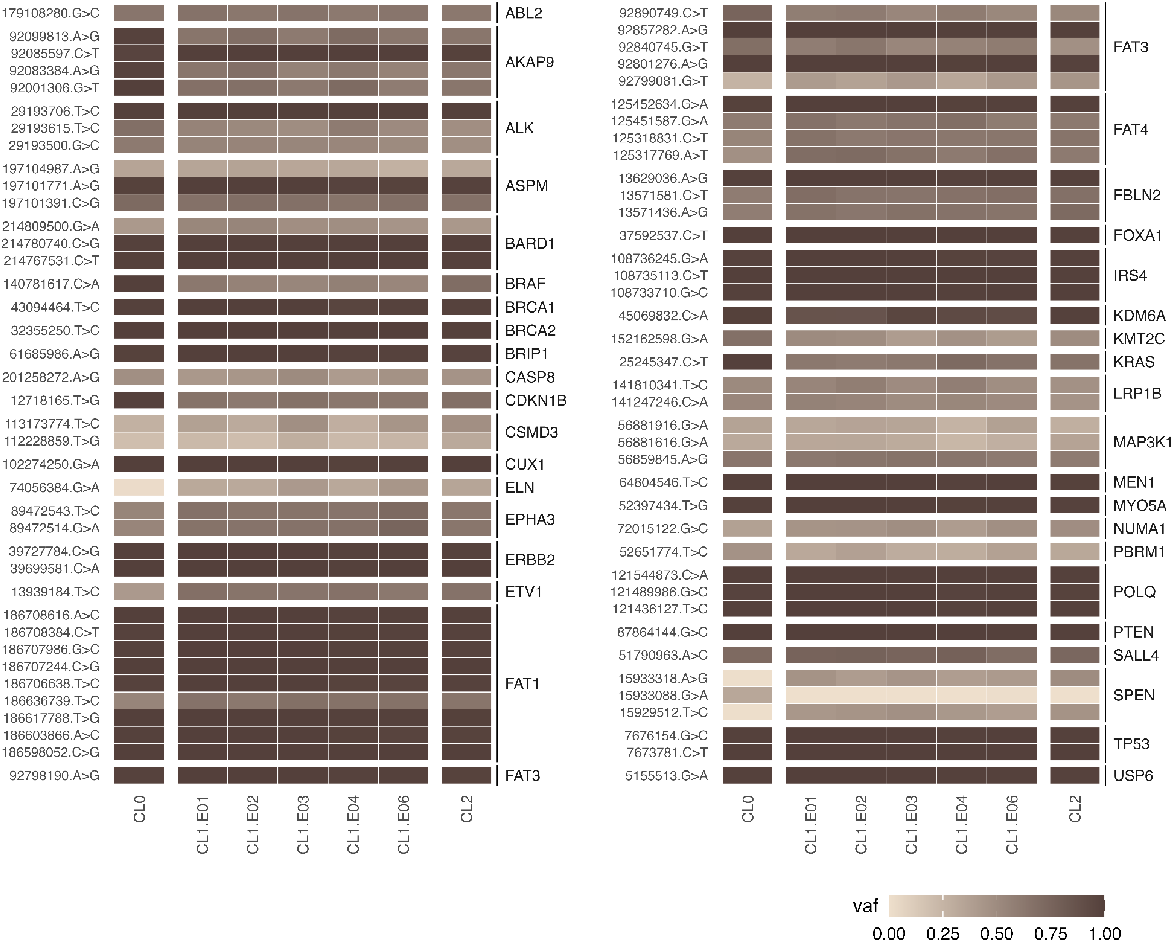
VAF of non-synonymous SNVs in BRCA driver genes. Each row represents a non-synonymous SNV in a specific gene (row facets) for each sample (columns). Mutations are labeled by their position (hg38 reference genome) and the amino acid change. VAF is depicted using a gradient color scale.

### Subline-specific mutational signatures revealed minimal variation

The mutational signatures showed minimal variation among sublines. When considering only mutations not shared by all the sublines, three single base substitution (SBS) signatures were common to all sublines (Figure 6A); SBS1 and SBS5 are clock-like signatures associated with endogenous mutational processes, while SBS54 is likely a sequencing artifact ^36^. Subline-specific signatures included SBS7a, related to UV light exposure ^37^, and SBS40, of unknown etiology, which were exclusively detected in CL0. SBS12, also of unknown etiology, was identified only in CL1. In CL2, we detected SBS32, associated with azathioprine treatment ^38^, and SBS37, of unknown etiology. Additionally, we found the UV-related doublet base substitution signature DBS1 ^39^ in CL0 (Figure 6B) and the unknown etiology signature DBS8 in CL1 and CL2. All sublines exhibited indel (ID) signatures ID1 and ID2 (Figure 6C), linked to replication slippage during DNA replication ^36^. These indel signatures are frequently observed in non-hypermutated tumors and are associated with SBS1 ^13^. When considering only mutations common to all sublines, we identified an additional signature, DBS3 (Figure S1), associated with polymerase epsilon exonuclease domain mutations (Alexandrov et al., 2020), not detected in the subline-specific analysis.

**Figure 6.**
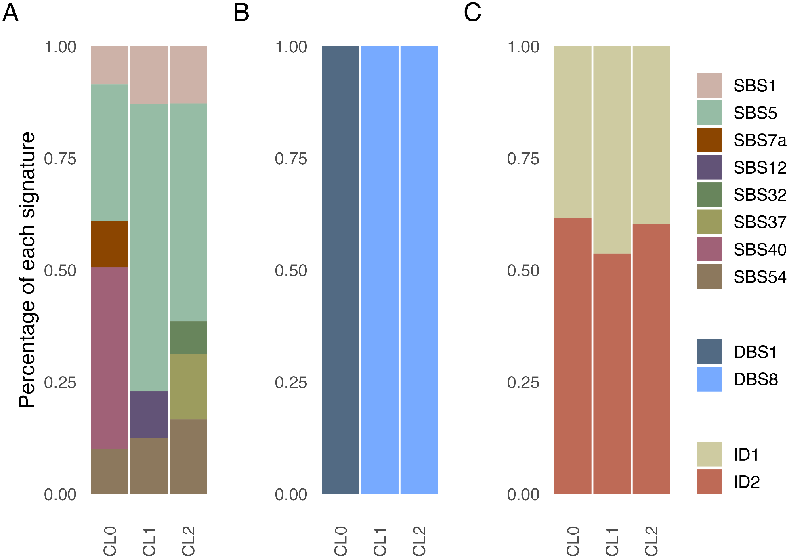
COSMIC subline-specific mutational signatures. Mutations shared across all sublines were excluded. (A) Single base substitution (SBS) signatures. (B) Doublet base substitution (DBS) signatures. (C) Small insertions and deletions (ID) signatures.

### Genomic similarity estimations depicted the relationship of MDA-MB-231 passages

Next, we investigated the genomic similarity among sublines. At the SNV level, all sublines were very similar (Figure 7A). In contrast, there were apparent differences regarding the CNA profiles (Figure 7B). CL0, the oldest subline, was the most distinct, both for SNVs and CNAs. CL1 passages (E01-E06) clustered together regarding their CNA profiles, but for SNVs, CL1.03 and CL1.06 clustered with CL2.

**Figure 7.**
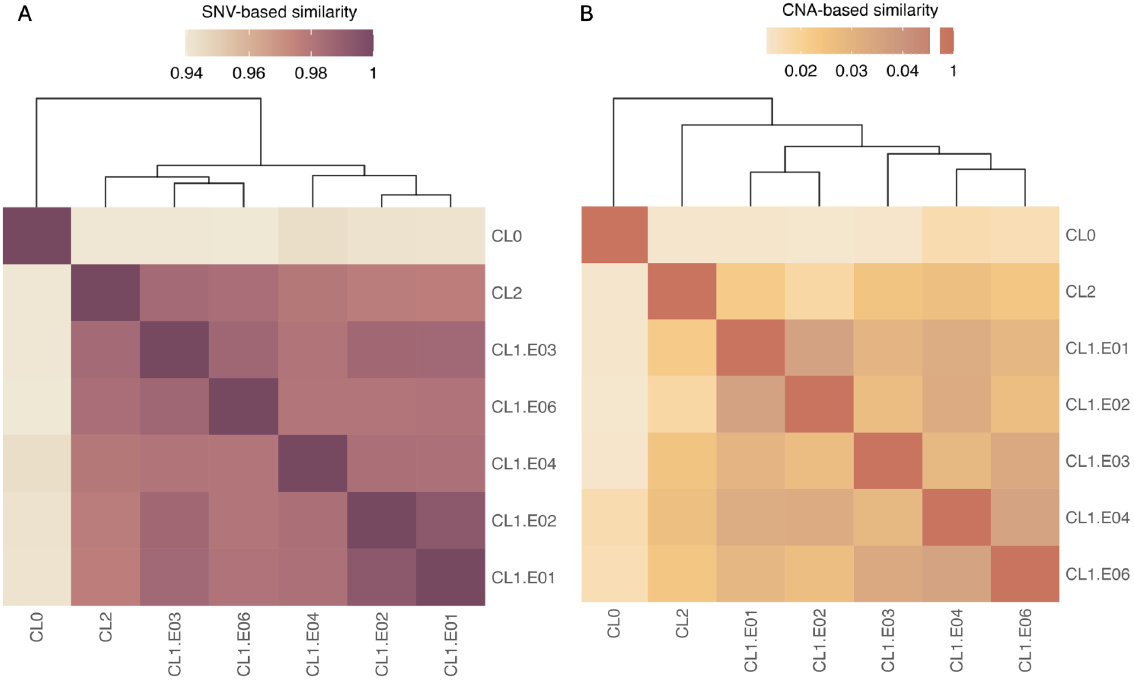
Genomic similarity measures. Comparative pairwise similarity estimations between sublines and passages. (A) SNV-based similarity coefficients were calculated using Jaccard. (B) CNA-based similarity measures are inferred with the *CNVMetrics* package using the log2-based option.

## DISCUSSION

In this study, we characterized the genomic heterogeneity among MDA-MB-231 sublines. We coined the term *intraline heterogeneity* (ILH) to describe the genomic variation observed both between sublines—cell lines purchased at distinct time points or from different providers—and across different culture passages produced in our laboratory.

Our karyotypic analysis suggests a near-triploidy chromosomal count, consistent with previous reports ^20,21^ and our CNA analysis. CNAs are a hallmark of breast cancer, particularly TNBC, where the most frequent alterations include gains of 1q (60%) and 8q; and losses of 8p ^40,41^. All three sublines exhibited 1q and 8q gains, while CL0 displayed a partial loss of 8p. Gene-level copy number analysis revealed significant heterogeneity across sublines, with shared breast cancer driver gene gains (e.g., *NOTCH1, TP53, ERBB2*) and losses (e.g., *JAK2* and *PTPRD*). CL0 harbored unique alterations, such as higher copy numbers in *PIK3CA* and *TBX3*. Apparently, all sublines exhibited considerable divergence at the CNA level, presumably driven by genomic instability. However, the CNA profiles were quite noisy, likely due to being inferred without a matched normal sample, and some of the observed differences might not reflect true biological variations.

In CL0, the oldest subline, three non-synonymous SNVs (*BRAF*-G464V, *KRAS*-G13D, and *CDKN1B*-V109G) showed an increased VAF. These variants have been reported as correlated with metastatic potential and worse prognosis in various cancers, including BRCA. Notably, *BRAF*-G464V and *KRAS*-G13D have been linked to tumor progression and metastatic potential in BRCA ^25,26^. Interestingly, in colorectal cancer models, a distinct amino acid change at the same position (*KRAS*-G13V) has been suggested to enhance metastatic potential ^42^. *CDKN1B* is a tumor suppressor gene infrequently mutated in cancer; however, when altered, it has been associated with increased tumor aggressiveness and poor clinical outcomes, particularly in spontaneous colitis-associated colorectal cancer ^43,44^. In the context of BRCA, *CDKN1B*-V109G has only been reported in non-TNBC ^32^, particularly in metastatic samples ^30,31,33^. The increased prevalence of this variant in metastatic samples proposes its potential as a biomarker. Further investigation is needed to determine whether this variant influences tumor behavior in TNBC and whether it could serve as a therapeutic target. Beyond these BRCA-associated variants, we identified three non-synonymous SNVs in *SPEN* with varying VAFs among sublines. Of particular interest, one of these (*SPEN*-R2283K) is reported for the first time in this study. Given that mutations and deletions in *SPEN* are related to tumor formation and progression ^45,46^, this novel variant could represent an unrecognized contributor to tumor evolution in MDA-MB-231 sublines. Taken together, we have found that all sublines acquired changes in allele frequencies in non-synonymous SNVs in BRCA driver genes throughout culture passages, which may potentially lead to significant differences in experimental outcomes.

Regarding mutational signatures, we found minimal variation among sublines. All sublines exhibited the clock-like signatures SBS1, SBS5, ID1, and ID2, which are associated with aging and are widely observed across cancer types ^36,47–50^. These signatures likely reflect the long-term accumulation of mutations rather than subline-specific selective pressures. Interestingly, CL0 harbored two unique subline-specific signatures, SBS7a and DBS1, linked to UV light exposure and frequently found together ^51,52^. The presence of these signatures calls attention to whether CL0 underwent environmental or culture-related stressors that contributed to their emergence. However, as MDA-MB-231 cells are not typically exposed to UV radiation, alternative mechanisms —such as oxidative stress or other mutagenic processes— may underlie these signatures. Further investigation is needed to determine the origin and potential functional impact of these mutational patterns in CL0.

In summary, we observe extensive ILH between MDA-MB-231 passages, often encompassing driver mutations that might result in phenotypic differences among sublines. These differences highlight the importance of considering ILH in experimental designs, as genomic differences between sublines may impact research outcomes. Future studies should assess the functional impact of these differences to improve the reproducibility and reliability of *in vitro* cancer research.

### Lead contact

Further information and requests for resources should be directed to and will be fulfilled by the corresponding author, David Posada.

## Acknowledgments

This work has been funded by the Spanish MICINN grant PID2019-106247GB-100 awarded to D.P. and by the PhD fellowship PRE2020-092269 awarded to N.V.R.

## Author contributions

Formal analysis, investigation, N.V.R.; cell line obtention and culture, C.F.S., and R.P.; cell culture for DNA extraction, doubling time, and sequencing library preparation, N.E.G., and M.P.; karyotype analysis, D.G.S, and J.J.P.; writing—original draft, N.V.R., J.M.A., and D.P; writing—review and editing, N.E.G., L.T..; conceptualization and supervision, J.M.A., and D.P.; funding acquisition, D.P.; all authors have read and reviewed the final version of this article.

## METHODS

### MDA-MB-231 cell lines

We analyzed three distinct commercial purchases of the MDA-MB-231 cell line (ATCC HTB26). This human triple-negative breast cancer cell line was established in 1973 from a pleural effusion of a 51-year-old Caucasian woman with metastatic breast cancer ^14^. We call each purchase of this cell line sublines CL0, CL1, and CL2. CL0, the oldest subline, was purchased at the American Type Culture Collection (ATCC) and modified *in situ* with lentiviral particles to express the Luciferase and the green fluorescent protein (GFP) ^53^ and has been maintained in our laboratory for over seven years. CL1 and CL2 were acquired four and two years ago, respectively, as dual-labeled luciferase/GFP variants (SL018; Genecopoeia, Inc, Rockville, MD, www.genecopoeia.com) from Tebu-Bio (www.tebubio.com) and AMSBIO (www.amsbio.com) correspondingly. Over the past four years, we produced five different culture passages of CL1 (CL1.E01, CL1.E02, CL1.E03, CL1.E04, and CL1.E06). Importantly, passage numbers (e.g., E01, E02, E03) do not necessarily correspond with chronology.

### Cell culture and DNA extraction

CL1 (MDA-MB-231-GFP) cells were cultured with DMEM w/o pyruvate medium (Gibco), enriched with 10% fetal bovine serum (FBS; ThermoFisher Scientific) and 1% penicillin/streptomycin (P/S; Gibco) under an atmosphere containing 5% CO_2_ at 37 ^º^C and a relative humidity level of 95%. We extracted genomic DNA (gDNA) of ∼ 1 million cells with AllPrep DNA/RNA/Protein mini kit (Qiagen) in 50 uL of the elution buffer. We estimated the concentration of gDNA with the dsDNA BR assay in a Qubit 3.0 (Invitrogen) fluorometer and the gDNA integrity using the Genomic DNA assay in a 2200 TapeStation platform (Agilent Technologies).

### Doubling time

We estimated the doubling time using the growth curve of CL1 (Figure S2). We seeded 125,000 cell triplicates in 6-well plates, counting for 10 days every 24 hours under the microscope. We used trypan blue 0.4 % (Gibco) and counted only unstained (live) cells. The media was changed every two days. The doubling time was calculated using the following formula:

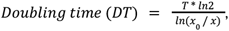

where *T* is the incubation time, *x*_0_ and *x* are the initial and final number of cells, respectively.

### Karyotype

We performed a cytogenetic analysis of CL1 following the standard protocol ^54^. Proliferating MDA-MB-231 cells were treated with 0.001% colchicine for 60 minutes, detached by scraping, and centrifuged (1,000g × 10 min). The cell pellet was subjected to a hypotonic treatment with warm 0.4% KCl (37°C × 30 min), followed by centrifugation (1,000g × 10 min) and fixation with a cold methanol and acetic acid (3:1) mixture. We prepared metaphase spreads using a standard splash technique, stained with DAPI (0.14 pg/ml), and mounted in an antifading medium (Vectashield, Vector). Finally, we captured cell images using a Nikon Eclipse-e800 fluorescence microscope equipped with a DS-Qi1Mc CCD camera controlled by NIS-Elements software (Nikon). We then processed the final images using Adobe Photoshop CS2 (Adobe Systems, San Jose, CA, USA).

### Whole exome sequencing

Sequencing libraries were constructed with the KAPA Hyper Prep and KAPA HyperExome kits (Roche Molecular Systems, Inc.) and sequenced at the Spanish National Center for Genomic Analysis (CNAG; http://www.cnag.crg.eu) on an Illumina NovaSeq 6000 (PE150) at a 1,000X sequencing depth.

### Preprocessing of sequencing data

We removed the *KAPA HyperExome* adapters using Cutadapt v.3.5 ^55^ and carried out quality control of all FASTQ samples with fastqQC v.0.12.1 ^56^ and MultiQC v.1.14 ^57^. Afterward, we mapped the trimmed reads to the human reference genome hg38 with BWA mem v.0.7.17 ^58^. Following the Genome Analysis Toolkit (GATK) Best Practices ^59,60^, we used Picard Tools v.2.25.5 (https://broadinstitute.github.io/picard/) SortSam and MarkDuplicates to sort the resulting SAM files and tag duplicates, respectively. Finally, we used GATK’s BaseRecalibrator and ApplyBQSR v.4.2.0 to recalibrate the base qualities.

### Variant calling

We identified single nucleotide variants (SNVs) and short insertions and deletions (indels) using HaplotypeCaller v.4.2.6.1 ^61^ in joint mode. Following GATK’s Best Practices, we applied standard filters using Variant Quality Score Recalibration (VQSR) v.4.2.6.1. In addition, we only kept biallelic variants, which were annotated using Annovar v.2020.06.07 ^62^.

We identified copy number alterations (CNAs) using Sequenza v.3.0.0 ^63^. Since we lack a matched normal sample, we used a healthy human dermal fibroblast (HDF) cell line processed under the same conditions as our MDA-MB-231 sublines. Sequenza was initially run under default parameters to obtain estimates of tumor purity and mean ploidy across sublines (Figure S3). Based on these results, considering that our samples comprise pure cell lines, we set the sample purity to 1. The mean ploidy was set to 2.7, calculated as the average of the ploidy estimates across all sublines. These parameter values were incorporated into Sequenza’s *baf*.*bayes* function to adjust for tumor-normal admixture and aneuploidy. This should enable more precise identification of CNAs by aligning observed sequencing data with the expected purity and ploidy scores.

### Mutational signatures

We inferred single base substitutions (SBS), doublet base substitutions (DBS), and small insertions and deletions (ID) mutational signatures (see https://cancer.sanger.ac.uk/signatures) for each subline using SigProfilerAssignment v.0.0.32 ^64^ in exome mode. To investigate shared and unique mutational processes, we constructed two datasets, one containing only mutations (SNVs and indels) not present in all sublines and another containing the remaining mutations shared by all sublines.

### Genomic similarity

We computed SNV-based pairwise similarities among samples using the Jaccard coefficient ^65^ formula as follows:

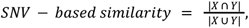

where *X* and *Y* are the two samples involved in the pairwise comparison. The coefficient is calculated as SNVs common to both samples divided by all the SNVs present in *X* or in *Y*.

For CNAs, we used CNVMetrics v.1.5.1 ^66^ to calculate pairwise similarities based on the *log*_2_ depth ratio, following the equation:

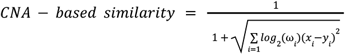

where *x*_*i*_ is the *log*_2_ ratio of segment *i* for sample *x, y*_*i*_ is the *log* ratio of segment *i* for sample *y*, and *ω*_*i*_ is the length in base pairs of the fixed segment *i*.

## SUPPLEMENTAL INFORMATION

**Table S1.** Sequencing metrics. Respectively, sample name (sample), mean depth (mean DP), percentage of reads mapped to human (hg38) (% human reads), and unmapped (% unmapped reads). We also report the percentage of the genome covered by at least 1 (% breadth 1X), 10 (% breadth 10X), and 100 (% breadth 100X) reads. All referred to the target exomic regions from the hg38 KAPA HyperExome kit.

| sample | mean DP | % human reads | % unmapped reads | % Breadth 1X | % Breadth 10X | % Breadth 100X |
| --- | --- | --- | --- | --- | --- | --- |
| CL0 | 303.45 | 99.80 | 0.20 | 99.68 | 99.64 | 99.40 |
| CL1.E1 | 763.12 | 99.81 | 0.19 | 99.69 | 99.66 | 99.50 |
| CL1.E2 | 507.44 | 99.77 | 0.22 | 99.69 | 99.66 | 99.42 |
| CL1.E3 | 1096.96 | 99.81 | 0.19 | 99.69 | 99.66 | 99.55 |
| CL1.E4 | 293.11 | 99.81 | 0.19 | 99.69 | 99.64 | 99.35 |
| CL1.E6 | 313.89 | 99.79 | 0.20 | 99.69 | 99.65 | 99.44 |
| CL2 | 507.28 | 99.81 | 0.19 | 99.74 | 99.67 | 99.54 |

**Table S2.** Non-synonymous SNVs identified in this study. Gene symbol, genomic position for the hg38 reference, amino acid (aa) change, ClinVar ^27^ aa change, aa change reported in COSMIC for MDA-MB-231 cell line, and verified status in COSMIC for MDA-MB-231 cell line ^13^.

| Gene symbol | Genomic position for hg38 | Amino acid change | ClinVar amino acid change | Reported for MDA-MB-231 in COSMIC | Verified in COSMIC |
| --- | --- | --- | --- | --- | --- |
| *ABL2* | chr1:179108280_G>C | - | - |  |  |
| *AKAP9* | chr7:92001306_G>T | M463I | p.Met463Ile |  |  |
| *AKAP9* | chr7:92083384_A>G | N2792S | p.Asn2792Ser |  |  |
| *AKAP9* | chr7:92085597_C>T | P2979S | p.Pro2979Ser |  |  |
| *AKAP9* | chr7:92099813_A>G | M3614V | p.Met3614Val |  |  |
| *ALK* | chr2:29193500_G>C | D1529E | p.Asp1529Glu |  |  |
| *ALK* | chr2:29193615_T>C | K1491R | p.Lys1491Arg |  |  |
| *ALK* | chr2:29193706_T>C | I1461V | p.Ile1461Val |  |  |
| *ASPM* | chr1:197101391_C>G | Q2620H | p.Gln2620His |  |  |
| *ASPM* | chr1:197101771_A>G | Y2494H | p.Tyr2494His |  |  |
| *ASPM* | chr1:197104987_A>G | - | - | S1422P | verified |
| *BARD1* | chr2:214767531_C>T | V507M | p.Val507Met |  |  |
| *BARD1* | chr2:214780740_C>G | R378S | p.Arg378Ser |  |  |
| *BARD1* | chr2:214809500_G>A | P24S | p.Pro24Ser |  |  |
| *BRAF* | chr7:140781617_C>A | G464V | p.Gly464Val | G464V | verified |
| *BRCA1* | chr17:43094464_T>C | Q356R | p.Gln356Arg |  |  |
| *BRCA2* | chr13:32355250_T>C | V2466A | p.Val2466Ala |  |  |
| *BRIP1* | chr17:61685986_A>G | S919P | p.Ser919Pro |  |  |
| *CASP8* | chr2:201258272_A>G | K14R | p.Lys14Arg |  |  |
| *CDKN1B* | chr12:12718165_T>G | V109G | p.Val109Gly |  |  |
| *CSMD3* | chr8:112228859_T>G | N3621H | p.Asn3621His |  |  |
| *CSMD3* | chr8:113173774_T>C | I219M | p.Ile219Met |  |  |
| *CUX1* | chr7:102274250_G>A | A464T | p.Ala464Thr |  |  |
| *ELN* | chr7:74056384_G>A | G422S | p.Gly422Ser |  |  |
| *EPHA3* | chr3:89472514_G>A | - | - |  |  |
| *EPHA3* | chr3:89472543_T>C | - | - |  |  |
| *ERBB2* | chr17:39699581_C>A | P8T | p.Pro8Thr |  |  |
| *ERBB2* | chr17:39727784_C>G | P1170A | p.Pro1170Ala |  |  |
| *ETV1* | chr7:13939184_T>C | - | - |  |  |
| *FAT1* | chr4:186598052_C>G | K4059N | p.Lys4059Asn |  |  |
| *FAT1* | chr4:186603866_A>C | S3554A | p.Ser3554Ala |  |  |
| *FAT1* | chr4:186617788_T>G | Q2933P | p.Gln2933Pro |  |  |
| *FAT1* | chr4:186636739_T>C | H1273R | p.His1273Arg |  |  |
| *FAT1* | chr4:186706638_T>C | R1064G | p.Arg1064Gly |  |  |
| *FAT1* | chr4:186707244_C>G | V862L | p.Val862Leu |  |  |
| *FAT1* | chr4:186707986_G>C | F614L | p.Phe614Leu |  |  |
| *FAT1* | chr4:186708384_C>T | V482I | p.Val482Ile |  |  |
| *FAT1* | chr4:186708616_A>C | S404R | p.Ser404Arg |  |  |
| *FAT3* | chr11:92798190_A>G | Q1726R | p.Gln1726Arg |  |  |
| *FAT3* | chr11:92799081_G>T | - | - | G1873V | unverified |
| *FAT3* | chr11:92801276_A>G | - | - |  |  |
| *FAT3* | chr11:92840745_G>T | V3518M | p.Val3518Met |  |  |
| *FAT3* | chr11:92857282_A>G | - | - |  |  |
| *FAT3* | chr11:92890749_C>T | - | - | S4319F | unverified |
| *FAT4* | chr4:125317769_A>T | Q453L | p.Gln453Leu |  |  |
| *FAT4* | chr4:125318831_C>T | A807V | p.Ala807Val |  |  |
| *FAT4* | chr4:125451587_G>A | G3526D | p.Gly3526Asp |  |  |
| *FAT4* | chr4:125452634_G>A | S3875N | p.Ser3875Asn |  |  |
| *FBLN2* | chr3:13571436_A>G | - | - |  |  |
| *FBLN2* | chr3:13571581_C>T | - | - |  |  |
| *FBLN2* | chr3:13629036_A>G | T901A | p.Thr901Ala |  |  |
| *FOXA1* | chr14:37592537_C>T | - | - |  |  |
| *IRS4* | chrX:108733710_G>C | H879D | p.His879Asp |  |  |
| *IRS4* | chrX:108735113_C>T | R411Q | p.Arg411Gln |  |  |
| *IRS4* | chrX:108736245_G>A | - | - |  |  |
| *KDM6A* | chrX:45069832_C>A | T778K | p.Thr778Lys |  |  |
| *KMT2C* | chr7:152162598_G>A | S3660L | p.Ser3660Leu |  |  |
| *KRAS* | chr12:25245347_C>T | G13D | p.Gly13Asp | G13D | verified |
| *LRP1B* | chr2:141247246_C>A | - | - | R191I | unverified |
| *LRP1B* | chr2:141810341_T>C | Q48R | p.Gln48Arg |  |  |
| *MAP3K1* | chr5:56859845_A>G | N255S | p.Asn255Ser |  |  |
| *MAP3K1* | chr5:56881616_G>A | D806N | p.Asp806Asn |  |  |
| *MAP3K1* | chr5:56881916_G>A | V906I | p.Val906Ile |  |  |
| *MEN1* | chr11:64804546_T>C | T541A | p.Thr541Ala |  |  |
| *MYO5A* | chr15:52397434_T>G | E362D | p.Glu362Asp |  |  |
| *NUMA1* | chr11:72015122_G>C | - | - |  |  |
| *PBRM1* | chr3:52651774_T>C | - | - | I228V | unverified |
| *POLQ* | chr3:121436127_T>C | Q2513R | p.Gln2513Arg |  |  |
| *POLQ* | chr3:121489986_G>C | T982R | p.Thr982Arg |  |  |
| *POLQ* | chr3:121544873_C>A | R66I | p.Arg66Ile |  |  |
| *PTEN* | chr10:87864144_G>C | S65C | p.Ser65Cys |  |  |
| *SALL4* | chr20:51790963_A>C | L507R | p.Leu507Arg |  |  |
| *SPEN* | chr1:15929512_T>C | L1091P | p.Leu1091Pro |  |  |
| *SPEN* | chr1:15933088_G>A | - | - |  |  |
| *SPEN* | chr1:15933318_A>G | N2360D | p.Asn2360Asp |  |  |
| *TP53* | chr17:7673781_C>T | R280K | p.Arg280Lys | R280K | verified |
| *TP53* | chr17:7676154_G>C | P72R | p.Pro72Arg |  |  |
| *USP6* | chr17:5155513_G>A | - | - |  |  |

**Figure S1.**
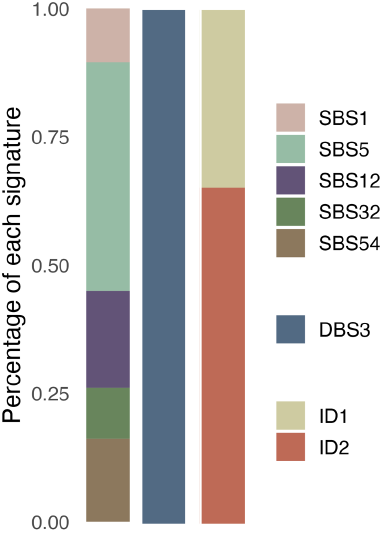
COSMIC mutational signatures for public mutations. We only consider mutations common to all sublines. Each bar corresponds to single base substitution (SBS) signatures, doublet base substitution (DBS) signatures, and small insertions and deletions (ID) signatures, respectively.

**Figure S2.**
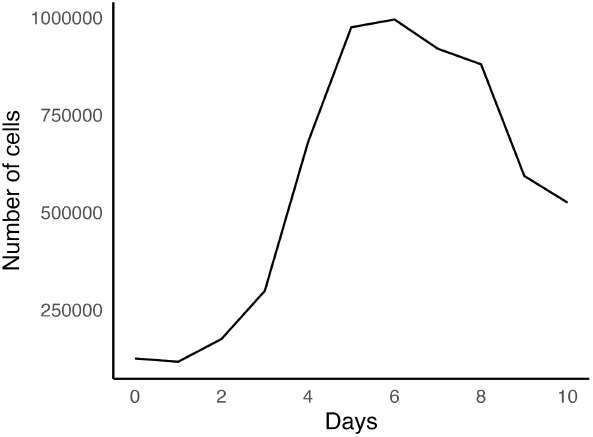
MDA-MB-231 growth curve. Mean number of viable cells (from triplicates) per day for the subline CL1.

**Figure S3.**
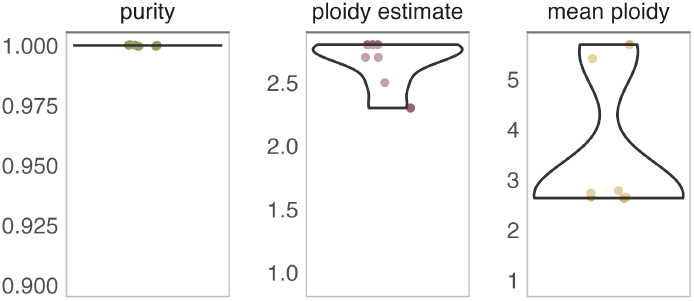
Purity, ploidy estimate, and mean ploidy estimates of Sequenza. Each dot corresponds to a subline.

## REFERENCES

Gillet, J.-P., Varma, S., and Gottesman, M.M. (2013). The clinical relevance of cancer cell lines. J. Natl. Cancer Inst. 105, 452–458. 10.1093/jnci/djt007.

Goodspeed, A., Heiser, L.M., Gray, J.W., and Costello, J.C. (2016). Tumor-derived cell lines as molecular models of cancer pharmacogenomics. Mol. Cancer Res. 14, 3–13. 10.1158/1541-7786.MCR-15-0189.

Kim, H.S., Sung, Y.-J., and Paik, S. (2015). Cancer cell line panels empower genomics-based discovery of precision cancer medicine. Yonsei Med. J. 56, 1186–1198. 10.3349/ymj.2015.56.5.1186.

Bose, B., and Bozdag, S. (2024). Identifying cell lines across pan-cancer to be used in preclinical research as a proxy for patient tumor samples. Commun. Biol. 7, 1101. 10.1038/s42003-024-06812-3.

Iorio, F., Knijnenburg, T.A., Vis, D.J., Bignell, G.R., Menden, M.P., Schubert, M., Aben, N., Gonçalves, E., Barthorpe, S., Lightfoot, H., et al. (2016). A landscape of pharmacogenomic interactions in cancer. Cell 166, 740–754. 10.1016/j.cell.2016.06.017.

Hynds, R.E., Vladimirou, E., and Janes, S.M. (2018). The secret lives of cancer cell lines. Dis. Model. Mech. 11. 10.1242/dmm.037366.

Li, J., Settivari, R.S., and LeBaron, M.J. (2019). Genetic instability of in vitro cell lines: Implications for genetic toxicity testing. Environ. Mol. Mutagen. 60, 559–562. 10.1002/em.22280.

Gisselsson, D., Lindgren, D., Mengelbier, L.H., øra, I., and Yeger, H. (2010). Genetic bottlenecks and the hazardous game of population reduction in cell line based research. Exp. Cell Res. 316, 3379–3386. 10.1016/j.yexcr.2010.07.010.

Ben-David, U., Siranosian, B., Ha, G., Tang, H., Oren, Y., Hinohara, K., Strathdee, C.A., Dempster, J., Lyons, N.J., Burns, R., et al. (2018). Genetic and transcriptional evolution alters cancer cell line drug response. Nature 560, 325–330. 10.1038/s41586-018-0409-3.

Kotsarenko, K., Vechtova, P., Lieskovska, J., Füssy, Z., Cabral-de-Mello, D.C., Rego, R.O.M., Alberdi, P., Collins, M., Bell-Sakyi, L., Sterba, J., et al. (2020). Karyotype changes in long-term cultured tick cell lines. Sci. Rep. 10, 13443. 10.1038/s41598-020-70330-5.

Ghandi, M., Huang, F.W., Jané-Valbuena, J., Kryukov, G.V., Lo, C.C., McDonald, E.R., 3rd, Barretina, J., Gelfand, E.T., Bielski, C.M., Li, H., et al. (2019). Next-generation characterization of the Cancer Cell Line Encyclopedia. Nature 569, 503–508. 10.1038/s41586-019-1186-3.

Forbes, S.A., Beare, D., Boutselakis, H., Bamford, S., Bindal, N., Tate, J., Cole, C.G., Ward, S., Dawson, E., Ponting, L., et al. (2017). COSMIC: somatic cancer genetics at high-resolution. Nucleic Acids Res. 45, D777–D783. 10.1093/nar/gkw1121.

Tate, J.G., Bamford, S., Jubb, H.C., Sondka, Z., Beare, D.M., Bindal, N., Boutselakis, H., Cole, C.G., Creatore, C., Dawson, E., et al. (2019). COSMIC: The Catalogue Of Somatic Mutations In Cancer. Nucleic Acids Res. 47, D941–D947. 10.1093/nar/gky1015.

Cailleau, R., Young, R., Olivé, M., and Reeves, W.J., Jr (1974). Breast tumor cell lines from pleural effusions. J. Natl. Cancer Inst. 53, 661–674. 10.1093/jnci/53.3.661.

Welsh, J. (2017). Modeling breast cancer in animals—considerations for prevention and treatment studies. In Animal Models for the Study of Human Disease, P. M. Conn, ed. (Academic Press), pp. 925–948. 10.1016/b978-0-12-809468-6.00035-8.

Chavez, K.J., Garimella, S.V., and Lipkowitz, S. (2010). Triple negative breast cancer cell lines: one tool in the search for better treatment of triple negative breast cancer. Breast Dis. 32, 35–48. 10.3233/BD-2010-0307.

Hollestelle, A., Nagel, J.H.A., Smid, M., Lam, S., Elstrodt, F., Wasielewski, M., Ng, S.S., French, P.J., Peeters, J.K., Rozendaal, M.J., et al. (2010). Distinct gene mutation profiles among luminal-type and basal-type breast cancer cell lines. Breast Cancer Res. Treat. 121, 53–64. 10.1007/s10549-009-0460-8.

Holliday, D.L., and Speirs, V. (2011). Choosing the right cell line for breast cancer research. Breast Cancer Res. 13, 215. 10.1186/bcr2889.

Harrell, J.C., Pfefferle, A.D., Zalles, N., Prat, A., Fan, C., Khramtsov, A., Olopade, O.I., Troester, M.A., Dudley, A.C., and Perou, C.M. (2014). Endothelial-like properties of claudin-low breast cancer cells promote tumor vascular permeability and metastasis. Clin. Exp. Metastasis 31, 33–45. 10.1007/s10585-013-9607-4.

Satya-Prakash, K.L., Pathak, S., Hsu, T.C., Olivé, M., and Cailleau, R. (1981). Cytogenetic analysis on eight human breast tumor cell lines: high frequencies of 1q, 11q and HeLa-like marker chromosomes. Cancer Genet. Cytogenet. 3, 61–73. 10.1016/0165-4608(81)90057-1.

Dugina, V., Shagieva, G., Novikova, M., Lavrushkina, S., Sokova, O., Kireev, I., and Kopnin, P. (2021). Impaired expression of cytoplasmic actins leads to chromosomal instability of MDA-MB-231 basal-like mammary gland cancer cell line. Molecules 26, 2151. 10.3390/molecules26082151.

Padilla-Nash, H.M., Barenboim-Stapleton, L., Difilippantonio, M.J., and Ried, T. (2006). Spectral karyotyping analysis of human and mouse chromosomes. Nat. Protoc. 1, 3129–3142. 10.1038/nprot.2006.358.

Sondka, Z., Bamford, S., Cole, C.G., Ward, S.A., Dunham, I., and Forbes, S.A. (2018). The COSMIC Cancer Gene Census: describing genetic dysfunction across all human cancers. Nat. Rev. Cancer 18, 696–705. 10.1038/s41568-018-0060-1.

Gonzalez-Perez, A., Perez-Llamas, C., Deu-Pons, J., Tamborero, D., Schroeder, M.P., Jene-Sanz, A., Santos, A., and Lopez-Bigas, N. (2013). IntOGen-mutations identifies cancer drivers across tumor types. Nat. Methods 10, 1081–1082. 10.1038/nmeth.2642.

Jacob, L.S., Vanharanta, S., Obenauf, A.C., Pirun, M., Viale, A., Socci, N.D., and Massagué, J. (2015). Metastatic competence can emerge with selection of preexisting oncogenic alleles without a need of new mutations. Cancer Res. 75, 3713–3719. 10.1158/0008-5472.CAN-15-0562.

Nagaria, T.S., Williams, J.L., Leduc, C., Squire, J.A., Greer, P.A., and Sangrar, W. (2013). Flavopiridol synergizes with sorafenib to induce cytotoxicity and potentiate antitumorigenic activity in EGFR/HER-2 and mutant RAS/RAF breast cancer model systems. Neoplasia 15, 939–951. 10.1593/neo.13804.

Landrum, M.J., Lee, J.M., Riley, G.R., Jang, W., Rubinstein, W.S., Church, D.M., and Maglott, D.R. (2014). ClinVar: public archive of relationships among sequence variation and human phenotype. Nucleic Acids Res. 42, D980–D985. 10.1093/nar/gkt1113.

Milne, R.L., Burwinkel, B., Michailidou, K., Arias-Perez, J.-I., Zamora, M.P., Menéndez-Rodríguez, P., Hardisson, D., Mendiola, M., González-Neira, A., Pita, G., et al. (2014). Common non-synonymous SNPs associated with breast cancer susceptibility: findings from the Breast Cancer Association Consortium. Hum. Mol. Genet. 23, 6096–6111. 10.1093/hmg/ddu311.

Frank, B., Wiestler, M., Kropp, S., Hemminki, K., Spurdle, A.B., Sutter, C., Wappenschmidt, B., Chen, X., Beesley, J., Hopper, J.L., et al. (2008). Association of a common AKAP9 variant with breast cancer risk: a collaborative analysis. J. Natl. Cancer Inst. 100, 437–442. 10.1093/jnci/djn037.

Naidu, R., Har, Y.C., and Taib, N.A.M. (2007). P27 V109G Polymorphism is associated with lymph node metastases but not with increased risk of breast cancer. J. Exp. Clin. Cancer Res. 26, 133–140.

Schöndorf, T., Eisele, L., Göhring, U.-J., Valter, M.M., Warm, M., Mallmann, P., Becker, M., Fechteler, R., Weisshaar, M.-P., and Hoopmann, M. (2004). The V109G polymorphism of the p27 gene CDKN1B indicates a worse outcome in node-negative breast cancer patients. Tumour Biol. 25, 306–312. 10.1159/000081396.

Viotto, D., Russo, F., Anania, I., Segatto, I., Rampioni Vinciguerra, G.L., Dall’Acqua, A., Bomben, R., Perin, T., Cusan, M., Schiappacassi, M., et al. (2021). CDKN1B mutation and copy number variation are associated with tumor aggressiveness in luminal breast cancer. J. Pathol. 253, 234–245. 10.1002/path.5584.

Rinaldi, J., Sokol, E.S., Hartmaier, R.J., Trabucco, S.E., Frampton, G.M., Goldberg, M.E., Albacker, L.A., Daemen, A., and Manning, G. (2020). The genomic landscape of metastatic breast cancer: Insights from 11,000 tumors. PLoS One 15, e0231999. 10.1371/journal.pone.0231999.

Richards, S., Aziz, N., Bale, S., Bick, D., Das, S., Gastier-Foster, J., Grody, W.W., Hegde, M., Lyon, E., Spector, E., et al. (2015). Standards and guidelines for the interpretation of sequence variants: a joint consensus recommendation of the American College of Medical Genetics and Genomics and the Association for Molecular Pathology. Genet. Med. 17, 405–424. 10.1038/gim.2015.30.

Ng, D., Johnston, J.J., Teer, J.K., Singh, L.N., Peller, L.C., Wynter, J.S., Lewis, K.L., Cooper, D.N., Stenson, P.D., Mullikin, J.C., et al. (2013). Interpreting secondary cardiac disease variants in an exome cohort. Circ. Cardiovasc. Genet. 6, 337–346. 10.1161/CIRCGENETICS.113.000039.

Alexandrov, L.B., Kim, J., Haradhvala, N.J., Huang, M.N., Tian Ng, A.W., Wu, Y., Boot, A., Covington, K.R., Gordenin, D.A., Bergstrom, E.N., et al. (2020). The repertoire of mutational signatures in human cancer. Nature 578, 94–101. 10.1038/s41586-020-1943-3.

Nik-Zainal, S., Kucab, J.E., Morganella, S., Glodzik, D., Alexandrov, L.B., Arlt, V.M., Weninger, A., Hollstein, M., Stratton, M.R., and Phillips, D.H. (2015). The genome as a record of environmental exposure. Mutagenesis 30, 763–770. 10.1093/mutage/gev073.

Inman, G.J., Wang, J., Nagano, A., Alexandrov, L.B., Purdie, K.J., Taylor, R.G., Sherwood, V., Thomson, J., Hogan, S., Spender, L.C., et al. (2018). The genomic landscape of cutaneous SCC reveals drivers and a novel azathioprine associated mutational signature. Nat. Commun. 9, 3667. 10.1038/s41467-018-06027-1.

Chen, J.-M., Férec, C., and Cooper, D.N. (2013). Patterns and mutational signatures of tandem base substitutions causing human inherited disease. Hum. Mutat. 34, 1119–1130. 10.1002/humu.22341.

Shahrouzi, P., Forouz, F., Mathelier, A., Kristensen, V.N., and Duijf, P.H.G. (2024). Copy number alterations: a catastrophic orchestration of the breast cancer genome. Trends Mol. Med. 30, 750–764. 10.1016/j.molmed.2024.04.017.

Watson, E.V., Lee, J.J.-K., Gulhan, D.C., Melloni, G.E.M., Venev, S.V., Magesh, R.Y., Frederick, A., Chiba, K., Wooten, E.C., Naxerova, K., et al. (2024). Chromosome evolution screens recapitulate tissue-specific tumor aneuploidy patterns. Nat. Genet. 56, 900–912. 10.1038/s41588-024-01665-2.

Paliogiannis, P., Cossu, A., Tanda, F., Palmieri, G., and Palomba, G. (2014). KRAS mutational concordance between primary and metastatic colorectal adenocarcinoma. Oncol. Lett. 8, 1422–1426. 10.3892/ol.2014.2411.

Choi, S.H., Barker, E.C., Gerber, K.J., Letterio, J.J., and Kim, B.-G. (2020). Loss of p27Kip1 leads to expansion of CD4+ effector memory T cells and accelerates colitis-associated colon cancer in mice with a T cell lineage restricted deletion of Smad4. Oncoimmunology 9, 1847832. 10.1080/2162402X.2020.1847832.

Huang, H., Qiu, D., Zhou, Z., Wu, B., Shao, L., Pu, Y., He, T., Wu, Y., Cui, D., and Zhong, F. (2023). A pan-cancer analysis for the oncogenic role of cyclin-dependent kinase inhibitor 1B in human cancers. Discov. Oncol. 14, 126. 10.1007/s12672-023-00746-8.

Légaré, S., Cavallone, L., Mamo, A., Chabot, C., Sirois, I., Magliocco, A., Klimowicz, A., Tonin, P.N., Buchanan, M., Keilty, D., et al. (2015). The estrogen receptor cofactor SPEN functions as a tumor suppressor and candidate biomarker of drug responsiveness in hormone-dependent breast cancers. Cancer Res. 75, 4351–4363. 10.1158/0008-5472.CAN-14-3475.

Légaré, S., Chabot, C., and Basik, M. (2017). SPEN, a new player in primary cilia formation and cell migration in breast cancer. Breast Cancer Res. 19. 10.1186/s13058-017-0897-3.

Alexandrov, L.B., Jones, P.H., Wedge, D.C., Sale, J.E., Campbell, P.J., Nik-Zainal, S., and Stratton, M.R. (2015). Clock-like mutational processes in human somatic cells. Nat. Genet. 47, 1402–1407. 10.1038/ng.3441.

Nik-Zainal, S., Alexandrov, L.B., Wedge, D.C., Van Loo, P., Greenman, C.D., Raine, K., Jones, D., Hinton, J., Marshall, J., Stebbings, L.A., et al. (2012). Mutational processes molding the genomes of 21 breast cancers. Cell 149, 979–993. 10.1016/j.cell.2012.04.024.

Alexandrov, L.B., Nik-Zainal, S., Wedge, D.C., Aparicio, S.A.J.R., Behjati, S., Biankin, A.V., Bignell, G.R., Bolli, N., Borg, A., Børresen-Dale, A.-L., et al. (2013). Signatures of mutational processes in human cancer. Nature 500, 415–421. 10.1038/nature12477.

Thatikonda, V., Islam, S.M.A., Autry, R.J., Jones, B.C., Gröbner, S.N., Warsow, G., Hutter, B., Huebschmann, D., Fröhling, S., Kool, M., et al. (2023). Comprehensive analysis of mutational signatures reveals distinct patterns and molecular processes across 27 pediatric cancers. Nat. Cancer 4, 276–289. 10.1038/s43018-022-00509-4.

Degasperi, A., Zou, X., Amarante, T.D., Martinez-Martinez, A., Koh, G.C.C., Dias, J.M.L., Heskin, L., Chmelova, L., Rinaldi, G., Wang, V.Y.W., et al. (2022). Substitution mutational signatures in whole-genome-sequenced cancers in the UK population. Science 376. 10.1126/science.abl9283.

Otlu, B., Díaz-Gay, M., Vermes, I., Bergstrom, E.N., Zhivagui, M., Barnes, M., and Alexandrov, L.B. (2023). Topography of mutational signatures in human cancer. Cell Rep. 42, 112930. 10.1016/j.celrep.2023.112930.

Viczián, A., and Kircher, S. (2010). Luciferase and green fluorescent protein reporter genes as tools to determine protein abundance and intracellular dynamics. In Plant Developmental Biology. Methods in Molecular Biology, L. Hennig and C. Köhler, eds. (Humana Press), pp. 293–312. 10.1007/978-1-60761-765-5_20.

Schempp, W., and Meer, B. (1983). Cytologic evidence for three human X-chromosomal segments escaping inactivation. Hum. Genet. 63, 171–174. 10.1007/bf00291539.

Martin, M. (2011). Cutadapt removes adapter sequences from high-throughput sequencing reads. EMBnet J. 17, 10. 10.14806/ej.17.1.200.

Andrews, S. (2010). FastQC: A Quality Control Tool for High Throughput Sequence Data. Preprint.

Ewels, P., Magnusson, M., Lundin, S., and Käller, M. (2016). MultiQC: summarize analysis results for multiple tools and samples in a single report. Bioinformatics 32, 3047–3048. 10.1093/bioinformatics/btw354.

Li, H. (2013). Aligning sequence reads, clone sequences and assembly contigs with BWA-MEM. arXiv [q-bio.GN]. 10.48550/ARXIV.1303.3997.

DePristo, M.A., Banks, E., Poplin, R., Garimella, K.V., Maguire, J.R., Hartl, C., Philippakis, A.A., del Angel, G., Rivas, M.A., Hanna, M., et al. (2011). A framework for variation discovery and genotyping using next-generation DNA sequencing data. Nat. Genet. 43, 491–498. 10.1038/ng.806.

Van der Auwera, G.A., Carneiro, M.O., Hartl, C., Poplin, R., Del Angel, G., Levy-Moonshine, A., Jordan, T., Shakir, K., Roazen, D., Thibault, J., et al. (2013). From FastQ data to high confidence variant calls: the Genome Analysis Toolkit best practices pipeline. Curr. Protoc. Bioinformatics 43, 11.10.1–11.10.33. 10.1002/0471250953.bi1110s43.

Poplin, R., Ruano-Rubio, V., DePristo, M.A., Fennell, T.J., Carneiro, M.O., Van der Auwera, G.A., Kling, D.E., Gauthier, L.D., Levy-Moonshine, A., Roazen, D., et al. (2017). Scaling accurate genetic variant discovery to tens of thousands of samples. bioRxiv. 10.1101/201178.

Wang, K., Li, M., and Hakonarson, H. (2010). ANNOVAR: Functional annotation of genetic variants from next-generation sequencing data. Nucleic Acids Research 38.

Favero, F., Joshi, T., Marquard, A.M., Birkbak, N.J., Krzystanek, M., Li, Q., Szallasi, Z., and Eklund, A.C. (2015). Sequenza: allele-specific copy number and mutation profiles from tumor sequencing data. Ann. Oncol. 26, 64–70. 10.1093/annonc/mdu479.

Díaz-Gay, M., Vangara, R., Barnes, M., Wang, X., Islam, S.M.A., Vermes, I., Duke, S., Narasimman, N.B., Yang, T., Jiang, Z., et al. (2023). Assigning mutational signatures to individual samples and individual somatic mutations with SigProfilerAssignment. Bioinformatics 39. 10.1093/bioinformatics/btad756.

Jaccard, P. (1901). Étude comparative de la distribution florale dans une portion des Alpes et du Jura. 10.5169/SEALS-266450.

Belleau, P., Deschênes, A., Beyaz, S., Tuveson, D.A., and Krasnitz, A. (2021). CNVMetrics package: Quantifying similarity between copy number profiles. 10.7490/F1000RESEARCH.1118704.1.

